# How can brain-like dynamics and phonetic generalization emerge in speech representations? A study using self-supervised models

**DOI:** 10.64898/2026.01.16.700011

**Authors:** Oli Danyi Liu, Hao Tang, Naomi H. Feldman, Sharon Goldwater

**Affiliations:** University of Edinburgh; University of Maryland College Park

**Keywords:** speech processing, neural representations, computational modeling

## Abstract

Speech representations in the human brain do not simply mirror the instantaneous speech signal; rather, they display several properties that are hypothesized to facilitate the integration of speech sounds into words. In particular, neural encodings of speech maintain information that has dissipated from the acoustics, and have also been argued to abstract over variability in how individual speech sounds are produced. Here, we investigate how such characteristics could arise. We introduce a computational framework that uses modern neural network models from speech technology to examine two factors in particular: the learning mechanism and the learning input. We find that self-supervised models trained without lexical or semantic feedback develop temporal dynamics similar to brain representations, regardless of whether they were trained on speech or non-speech audio. In contrast, only models trained on speech learn to abstract over variability due to word position and phonetic context. Overall, our results suggest that domain-general learning mechanisms can lead to several important properties of speech representations, but in some cases require domain-specific input in order to do so.

**Significance Statement:** Understanding speech often feels effortless, but in fact mapping speech sounds into words involves complex computation. Experimental neuroscience has identified key properties in brain signals that may support this computation, but *why* and *how* these properties arise is still unclear. We examined these properties in computational models and found that they occurred in models that weren’t given lexical or semantic feedback, but were trained to predict the acoustics of the speech signal. This suggests such properties can develop from domain-general learning combined with domain-specific input. More-over, some properties even arose in models that were trained on non-speech audio. Overall, our work illustrates how computational modeling can help reveal the conditions under which neural properties emerge.

---

**S**peech recognition involves mapping continuous acoustic signals to a sequence of discrete linguistic units. The smallest such unit is a phone, each lasting around 80 ms in the acoustic stream. Yet humans experience speech as words, which need to be composed by tracking and integrating sequentially presented phones, each of which is perceived only transiently in the speech signal. The same set of phones can form distinct words, e.g. *tap, apt* and *pat*, so the relative ordering of the phones in a sequence also needs to be maintained. Furthermore, the acoustic realization of any particular phone can vary considerably due to *co-articulation*: that is, because speech is produced using continuous articulatory gestures, the articulation of each phone affects that of its neighbors, causing acoustic variability. This variability needs to be abstracted away in order for phones to be identified. Understanding how the brain solves these computational problems is essential for uncovering the mechanisms underlying human speech processing.

Recent work has studied how phonetic information is encoded in the brain (1–4), and identified several properties that could help tackle the transience and variability of speech signals. First, the brain maintains a representation of each phone for an extended period. Studies of both EEG and MEG recordings have shown that the encoding^*^ of a phone persists until long after its dissipation in the acoustics (3, 5), which allows successive phones to be maintained simultaneously before they are integrated. Second, the neural encoding pattern of each phone evolves over time (5). This kind of *dynamic code* jointly encodes the identity of a phone and how long ago it occurred, which provides a way to track the relative ordering between successive phones. Third, the encoding pattern of a phone is partially independent of its phonetic context (6) or its position within a word (5). This finding points towards but does not fully establish the presence of invariant representations of phones, which are crucial for mapping variable acoustics to discrete linguistic units. Although the discovery of these properties has begun to shed light on the nature of speech representations in the brain and how they might support speech processing, we still lack an account for how or why these neural phenomena arise. For instance, we do not know whether feedback from top-down information (such as explicit recognition of word forms or meanings) is needed to learn such representations. Although behavioral experiments have investigated how lexical knowledge affects phonetic categorization (7–9), they are limited in their capacity to probe how top-down feedback shapes pre-lexical representations during processing (10, 11), and even more so for learning. Experimental methods are also not well suited for determining whether these properties arise from general auditory learning or speech-specific learning. Since the transient nature of the perceptual input is not unique to speech but shared across most types of auditory signals, any representational dynamics that compensate for this transience could be domain general.

In this work, we use representation learning models from speech technology to investigate the learning mechanism and domain specificity underlying the neural phenomena found in humans processing continuous speech. We focus on the three key properties of phonetic encoding that were described above, namely maintenance over an extended period, dynamic encoding, and context invariance. We ask whether these properties manifest in the representations learned by three different models. All three models are based on a self-supervised learning framework, but differ in the implementation of the learning mechanism and/or the nature of their learning material. By examining whether each model displays the key properties found in brain representations, we can identify the constraints or conditions required for each property to arise.

## Simulating brain studies to analyze the conditions under which representational properties can arise

We use state-of-the-art self-supervised learning (SSL) models to simulate and extend results from brain recordings of human listeners. These artificial neural network models learn to map acoustic waveforms to a sequence of high dimensional vectors, with each vector encoding the model’s representation at a particular timestep of the input. In this way, the model representations can be treated as analogous to the brain recordings we simulate, which also consist of high-dimensional timeseries.

The SSL speech models used in our study are trained on raw, unlabeled speech without any explicit linguistic supervision. Nevertheless, such models have been shown to encode a wide range of linguistic and speaker-related information (12–16). Empirically, they prove more effective than traditional acoustic features when used for engineering tasks like automatic speech recognition.

From a scientific perspective, SSL models have also demonstrated potential as models of speech processing in humans. Although they may not capture all relevant aspects of speech processing (17), there is evidence that they exhibit some behavioral patterns resembling those of human listeners (18–20) and that they account for variance in neural responses of the auditory cortex ((21–28); but see (29, 30)). Our goal is not to adjudicate the models’ overall similarity to human processing, but rather, to replicate particular representational patterns that have been found in human neural recordings and to examine under what conditions those patterns arise. In our simulations, we compared three SSL models—*CPC-speech, CPC-audio*, and *Wav2Vec2* —to examine how specific neural dynamics are shaped by different types of learning mechanisms and learning materials. CPC-speech (31) and CPC-audio (32) are both implementations of the Contrastive Predictive Coding (CPC) framework (33). As illustrated in Figure 1(a), they are trained to learn representations that are useful for predicting acoustics that come later. Wav2Vec2 (34), in contrast, learns representations that can be used to reconstruct masked segments of speech by drawing on both preceding and following context within the same utterance.^†^ To test domain specificity, we contrasted CPC-speech and CPC-audio, which differ in their learning materials. CPC-audio was only exposed to audio scenes, including animal vocalization and environmental sounds, whereas CPC-speech and Wav2Vec2 were trained on English audio books.

**Fig 1.**
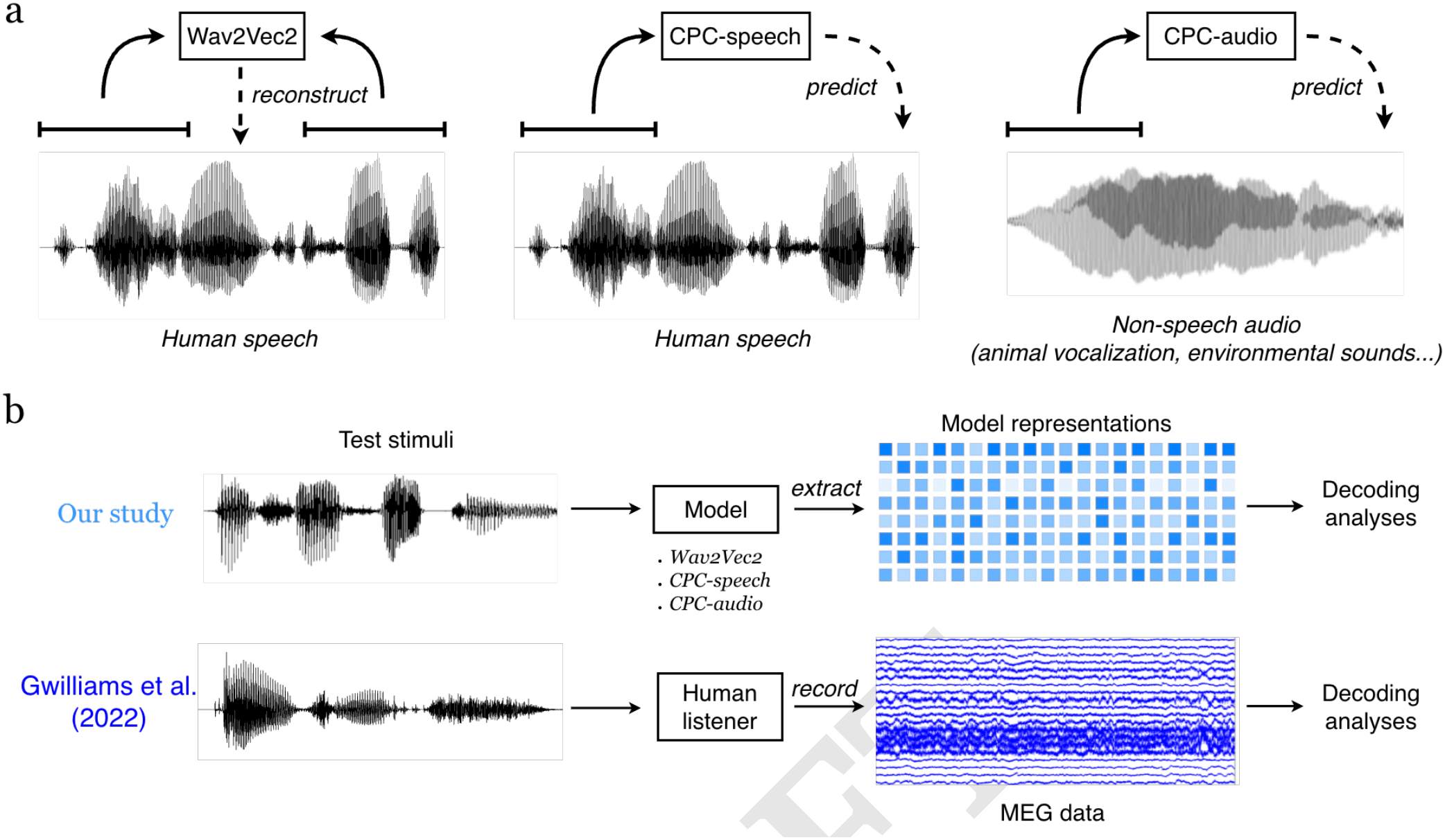
Experimental setup. (a) We considered three self-supervised learning models: Wav2Vec2 and CPC-speech differ in terms of learning mechanism — Wav2Vec2 was trained to reconstruct a part of an utterance given surrounding context while CPC was trained to predict upcoming acoustics based on past context in the same utterance — but are both trained on speech; CPC-audio shared the same learning mechanism as CPC-speech, but was trained on non-speech audio. (b) For each trained model, we fed in the same test stimuli (*speech*), extracted model representations, and applied decoding analyses analogous to those used in Gwilliams et al.’s analyses of MEG recordings from human listeners.

As shown in Figure 1b, we applied *decoding analyses* to representations extracted from each model. Such analyses are commonly used in cognitive neuroscience to study brain recordings. These analyses use machine learning-based classifiers (decoders) to identify fine-grained neural patterns associated with a target concept, which is usually a specific aspect of the perceptual stimuli, such as an acoustic or visual feature (35). Here, we follow Gwilliams et al. (5) in using *time-resolved* decoding, which can reveal not only whether the target information is present but also its temporal dynamics in the neural representations (36, 37). This type of analysis involves training a series of classifiers, each trying to identify the target based on a single time slice of the brain data. Mapping out the classifier accuracies across time reveals the time course of the neural encoding of interest. Time-resolved decoding is particularly well-suited for investigating time-sensitive processes such as speech processing, and especially when applied to neuro-imaging data with high temporal resolution. Furthermore, the extent to which the trained classifiers generalize across time, or from one condition to another, can provide insights into how the target is encoded over time and how consistent its neural encoding is across conditions.

For the decoders in our study, we trained multiclass logistic regression models to identify the phoneme category associated with each representation vector (henceforth *frame*), and evaluated their accuracy on a held-out test set. We consider phonetic information to be encoded if the decoding accuracy was higher than the majority class baseline (most common phoneme label). Phone boundaries and labels used in this process were obtained via force-aligning acoustics to the audio transcript using an acoustic model. As a reference, we also applied the same analyses on acoustic features (log Mel spectrograms) extracted from the same speech stimuli, in order to determine whether each property is already present in the acoustics.

## Results

### All three models maintain persistent phonetic encoding

We first examined the time window during which the phoneme category of a phone can be decoded from model representations or acoustic features. While the average duration of a phone is around 80 ms, neural recordings of human listeners suggest that phonetic features and/or phoneme identity are decodable from the brain for around 200–400 ms, with a latency of 10–50 ms after phone onset (3, 5). Such a decodable window extends well beyond the physical duration of a phone, implying that information about a phone is maintained even after it is no longer present in the acoustic signal. There is also evidence that SSL speech model representations encode contextual information over a considerably longer time window than the duration of a single phone (38, 39).

To evaluate the duration of phone decodability in our models, we considered a time window of 1200 ms centered around the onset of each phone sample. This window corresponds to 120 samples of CPC representations (which have a sample rate of 100 Hz). A separate decoder is trained for each of the timesteps to determine to what extent phones are decodable up to 600 ms before or 600 ms after their onset. For example, the −50 ms decoder for CPC is trained on frames that occur precisely 50 ms prior to a phone boundary, and must predict the phoneme label of the phone that starts at that boundary.

Before presenting the results, we first note that when decoding a phone, the decoder model may be accessing up to three different types of information: (1) *acoustic information* about the phone, including from co-articulation, which is presumably available for only a short time around the phone itself; (2) *sequential dependencies*, which give clues to what the preceding or following phone(s) might be, based on the current acoustics (for example, if the acoustic information available at −150 ms is most consistent with the sequence [st], then the phone starting at 0 ms is likely to be a vowel or [r]); and (3) *maintenance* of phonetic information in the model representations, i.e., representations that go beyond sequential dependencies to directly represent what *actually* happened at a distant point in the input, rather than what is *statistically likely* to happen. We are most interested in this third type of information, since its presence would simulate the type of maintenance found in the human brain data. Note that the CPC models (like humans) can only maintain information from previous parts of the utterance, while the Wav2vec2 model can (in principle) maintain information from both past and future, since it observes the entire utterance at once.

Figure 2 presents the results. Even for acoustic features, the decodable window extends beyond the phone boundaries, due to a combination of acoustic information and sequential dependencies. However, the models’ decodable windows are two to three times longer. Some of this extended window is also due to sequential dependencies, which we can see by the fact that phones are decodable from the CPC model representations considerably before their onset—i.e., before the models have heard any acoustic evidence of the phone. Moreover, the CPC-speech model, which has a chance to learn about speech-specific dependencies, has a longer pre-onset window of decodability than CPC-audio.

**Fig 2.**
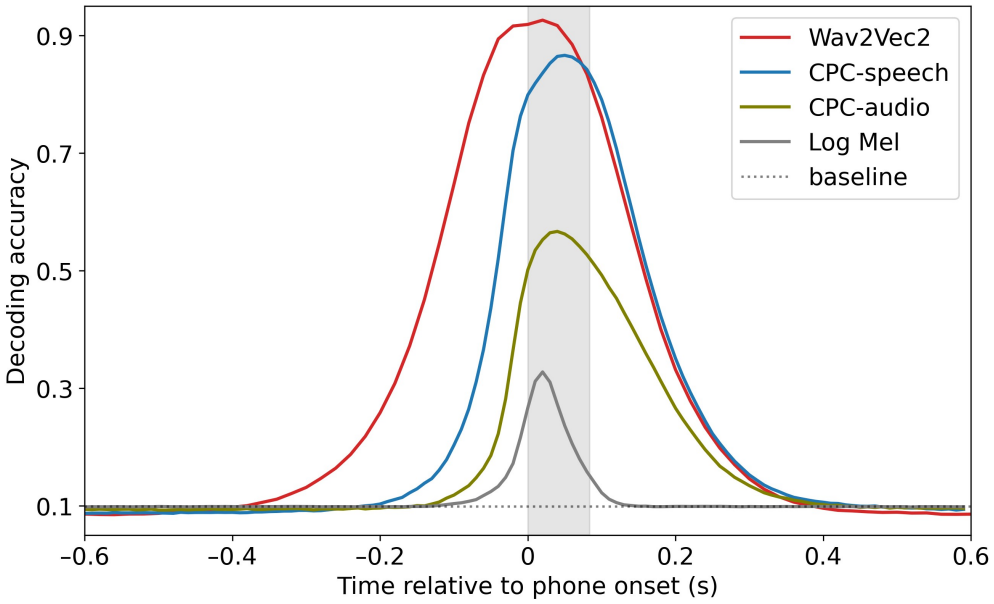
Decoding performance of time-resolved decoders within a 1400 ms sliding window centered on the onset of each phone. All three models we tested allow the phoneme identity to be decoded before and long after it is decodable in the acoustic features. Approximate decodable windows (relative to phone onset): CPC-speech: −200–400 ms; Wav2Vec2: −400–400 ms; CPC-audio: −130–400 ms; log Mel acoustic features: −110–120 ms. The dashed horizontal line shows the the majority class baseline (by predicting the most common phoneme label).

Crucially though, our results also show evidence of maintenance. In particular, the CPC models support decoding for much longer after phone offset than before phone onset. In addition, the Wav2Vec2 model, which has access to all input at once, shows a much longer pre-onset decodable window than the CPC models, suggesting that its representations are actually maintaining (i.e., directly representing) information about the upcoming phone(s), rather than just predicting which phone(s) are likely. Additional simulations provide further evidence that the asymmetric pattern of decodability in the CPC models cannot be explained by stimulus dependencies alone (see SI Text and Fig. S1, and (40) for further discussion of the general issue).

Finally, as compared to the log Mel spectrograms, model representations also support markedly higher decoding accuracy, indicating greater linear separability of phonemes in the models’ representation spaces. In particular, the audio model seems to have learned representations that are helpful for distinguishing between phoneme categories through general auditory structure alone, as its maximum decoding accuracy sits above the acoustic features. This suggests learning auditory structure alone can promote categorization of phones, although it is not as effective as learning from the structure of speech. While some of this gain might be explained by the higher dimensionality of the model representations, we also tested dimension-reduced representations that matched the dimensionality of acoustic features, and found that they still yielded wider decodable window and higher decoding accuracy (see SI Text and Fig. S2).

Overall, all three models maintained information about a phone for substantially longer than it is present in the acoustic signals, at a timescale that resembles that of neural encoding in humans. Notably, this property also emerged in CPC-audio, which was trained exclusively on non-speech sounds, and yet maintained information about a phone for nearly as long as CPC-speech and Wav2Vec2. This suggests that self-supervised learning can give rise to the sustained encoding found in brain recordings, without top-down feedback or domain-specific input.

### All three models jointly encode temporal and phonetic information with evolving representations

In the first simulation, we found that each phone is encoded for much longer in model representations than in acoustic features, which suggests explicit encoding of multiple successive phones simultaneously. This raises a question: how does a representation store information about successive phones while also preserving their order? One possibility is that the representation incorporates additional information about each phone. This information could indicate when a phone occurred (temporal distance from the current frame), where it occurred (its position within a word), or which other phones were around it (its phonetic context). If the encoding of a phone is conditioned on one or more types of such information, it would be possible to infer the phone’s relative order with respect to the other phones maintained simultaneously.

In this simulation, we focus on whether the encoding of a phone is conditioned on or independent of how long ago it occurred. For brains, Gwilliams et al. found evidence for temporally dependent encoding using temporal generalization (TG) analysis (41). We examined whether a similar phenomenon is present in the models, also through TG analyses. After training time-resolved decoders as in the previous simulation, we evaluated temporal generalization by taking each decoder—trained on a single timestep—and applying it to *all* the timesteps, including ones it hasn’t seen during training. Since each decoder has learned one most informative activation pattern for identifying the phonetic category at a certain moment (e.g. at phone onset), the extent to which each decoder generalizes to other timesteps (e.g. 50 ms into a phone) reflects whether that specific pattern persists till then.

To facilitate comparison with Gwilliams et al.’s results, we followed their approach by applying TG analysis to phone samples grouped into eight sets according to their position in the word: four positions moving forward from the first phone (p1–p4) and four moving backward from the last phone (p-1– p-4). Each TG analysis produces a matrix with dimension (training timestep) *×* (testing timestep), which is visualized as a contour plot, producing a figure that reflects the time course over which encoding of phones unfolds within a word, as shown in Figure 3. This allows us to pick a specific time and tell, on average, which phones are encoded at this time. For example, at 100 ms after word onset (which is on average during p2, as indicated by the red-orange bar on the x-axis), only p2 can be decoded from the log Mel spectrograms with accuracy above the threshold of 0.2. In contrast, for all three models (like the brain), multiple phones are decodable at this same 100 ms time point with an accuracy over 0.4: p1 and p2 in CPC-audio; p1–p3 in CPC-speech; and p1–p4 in Wav2Vec2.

**Fig 3.**
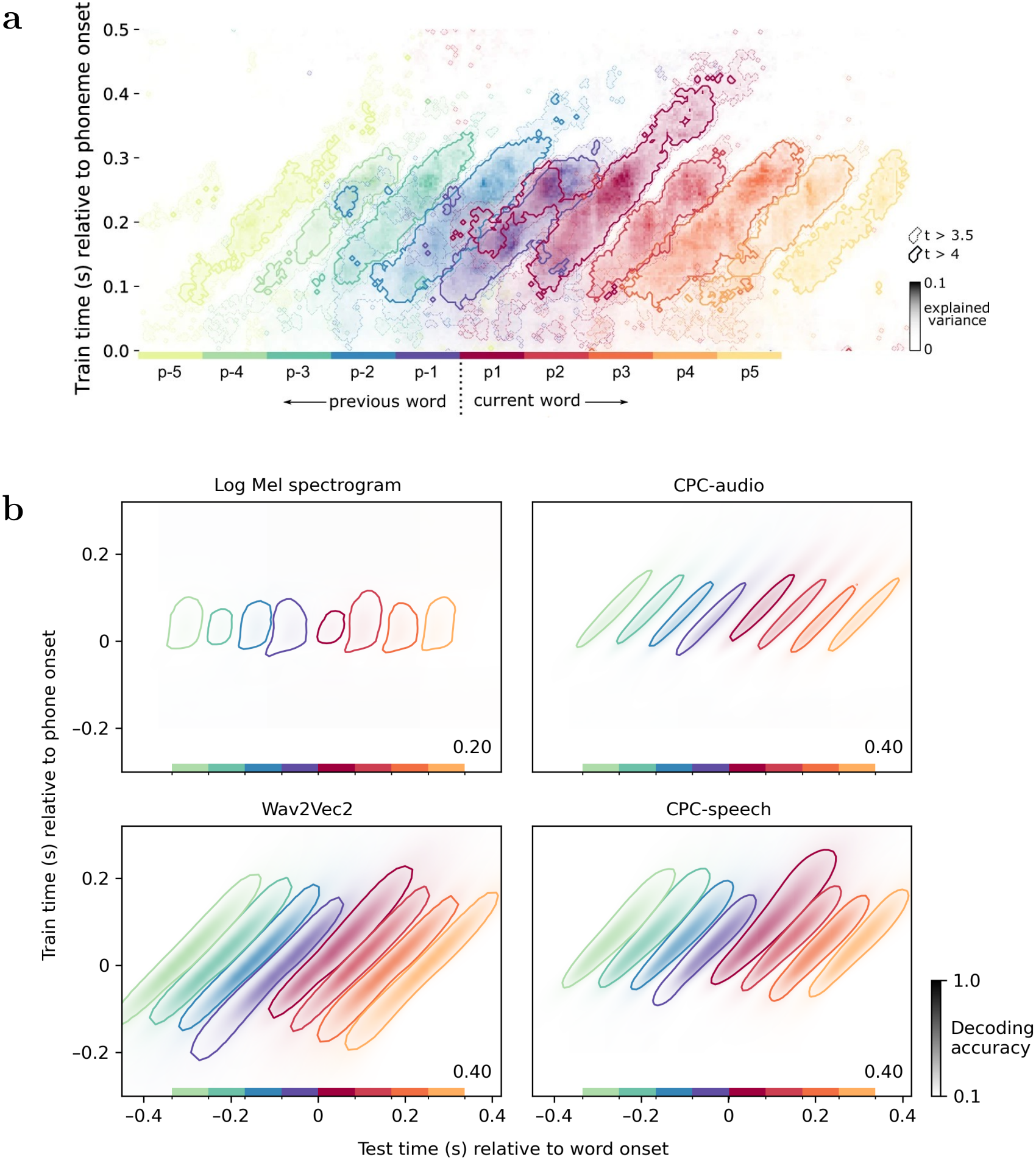
Temporal generalization results for (a) MEG recordings (reproduced from Gwilliams et al. (5)) and (b) acoustic features and model representations. Colors indicate phone positions within a word, with p1 (red) through p4 (yellow-orange) moving forward from the first phone and p-1 (purple) through p-4 (light green) moving backward from the last phone. The results for each phone position are shifted along the x-axis by the average phone duration (also visualized as colored bars on x-axis). Contours in (a) represent a t-value threshold of 4 (darker lines) and 3.5 (lighter lines), and in (b) show decoding accuracy thresholds of 0.4 (for the three models) or 0.2 (for acoustic features). The threshold chosen for the models, which occurs roughly 200 ms after phone onset for both Wav2Vec2 and CPC-speech, represents a clear improvement over the baseline but still far below maximum decoder accuracy. Both brain and model representations show long diagonal TG contours, indicating that encoding patterns change during the extended decodable window. For log Mel spectrograms, the diagonal and horizontal axes of each contour have similar lengths, indicating each phone is encoded with a pattern that is largely stable over time.

Figure 3 also reveals striking similarities in the shapes of the TG contours between the brain and all three models. In particular, the diagonal axis of each contour, which signifies the period that a phone is *decodable* from the representations, is much longer than any of its horizontal sections, which show the duration that each *neural pattern* persists. For example, in CPC-speech, with an accuracy threshold of 0.4, a phone is decodable for more than 200 ms on average, yet individual encoding patterns typically persist for no more than 100 ms—except for certain patterns associated with word-initial phones (p1). In constrast, the TG contours of acoustic features exhibit more temporally stable encoding patterns.

To confirm that these differences are due to learning, rather than to differences in dimensionality, we also tested reduced-dimensionality model representations and representations from untrained (randomly initialized) models (see Figure S4). The untrained models had very poor decoding accuracy but with hints of diagonal TG structure, suggesting that this structure can begin to emerge from random projections in a sufficiently high dimensional space. However, the reduced-dimensional model representations displayed both high accuracy and strong diagonal TG structure. We therefore conclude that learning, from either speech or non-speech audio, is more important than high dimensionality.

Given that these models are learning to maintain acoustic/phonetic information over time, we next asked what forms of maintenance could give rise to the TG patterns we observed. To do so, we examined two other representations: time-smoothed acoustic features and stacked acoustic features. Both retain information across a temporal window, but do so in different ways. The former additively combine log Mel features across time, representing each timestep as a weighted sum of the acoustic features from the current and several previous timesteps. The latter are constructed to maintain acoustic history in distinct dimensions, using the same number of dimensions as the representations from CPC-speech. Specifically, the feature vector for each timestep is created by concatenating the 40-dimensional log Mel feature vector of the current frame with those of the 12 preceding frames in temporal order, resulting in a vector of 13 *×* 40 = 520 dimensions. Thus, in these stacked features, information about the past is contained within each representation, but moves through different dimensions over time.

Results from this comparison (Figure S4) showed that Log Mel spectrograms still showed stable representations when smoothed over time, but exhibited evolving patterns when stacked across time. That is, diagonal TG patterns can arise from a representation in which acoustic context is maintained veridically, with different dimensions encoding information from different points in time. This suggests that the models could be implementing a similar approach, using a low-dimensional pattern to represent each input timestep, and shifting that same pattern across dimensions of the full representation space over time. However, the model representations are more efficient than the stacked log Mels, in the sense that with the same number of dimensions, they have higher decoding accuracy and a longer decodable window (diagonal axis). In fact, the decodable window in the stacked features is more similar to that of the reduced-dimensional model features. Thus, this type of veridical maintenance is not a sufficient explanation for what the models are learning. Finally, this simulation suggests a possible explanation for why the untrained models showed hints of diagonal structure: for any input timestep, random weights will incorporate some of the input information into internal representations for surrounding timesteps, and which internal dimensions that information can be found in will be different for different relative timesteps.

Finally, we identified a similarity between TG patterns of brains and CPC-speech that is not found in either Wav2Vec2 or CPC-audio. From both MEG signals and CPC-speech representations, word-initial phones (p1) remain decodable for a considerably longer period than phones in other positions (as indicated by their longer diagonal axis). While p3 and p4 had fewer training samples than p1, which could explain their narrower decodable windows, p1 and p2 had the same number of training samples. Additionally, the distribution of phonetic categories at p2 even had a lower entropy than at p1 (p1:3.11 bits, p2:3.03 bits), despite p1 being slightly longer in average duration (p1: 84.8ms, p2: 82.3ms). Again, the longer retention of word-initial phones is consistent with behavioral results showing their special role in word recognition (42). The absence of this pattern in Wav2Vec2 is likely because the model utilizes both preceding and subsequent contexts, while CPC relies solely on past context to predict upcoming acoustics, consequently develop a stronger reliance on word-initial phones. This suggests that the special significance of word-initial phones may not result from acoustic prominence, as they are not decodable for longer in the other two models or in acoustic features, but could be driven by the inherent directionality of speech processing. In this light, the use of bidirectional context in Wav2Vec2, while beneficial for engineering tasks, could limit its validity for modeling the representations underlying human speech perception.

Overall our results indicate that the joint encoding of phonetic content and temporal information can arise through self-supervised learning, including from non-speech audio. While this property can be useful for integrating phones into words, it is not necessarily shaped by the specific task demand of identifying words from speech.

### Learning from speech can induce positional and context invariance

So far we have shown that for each of the successive phones being maintained simultaneously, the phonetic content is jointly encoded with temporal information, allowing the relative ordering of the phones to be inferred. In theory, therefore, phone sequences could be integrated into words even if their representations abstract away from other sources of information about their ordering, such as their position in a word or the phonetic identities of neighboring sounds.

Here we examine whether such abstraction occurs, and under what conditions. Abstraction is central to many phonological theories, but it is not yet known what type of learning process enables listeners to abstract across contexts. A phone’s acoustic realization is heavily influenced by factors such as the manner and place of articulation of neighboring phones (43–45), meaning that the speech signal itself contains information about context. As a result, models of phonetic category learning have struggled to abstract across within-category contextual variability when trained directly on the acoustics (46, 47). On the other hand, there is evidence from both adults and infants that decoders trained on neural representations of the speech signal can partially generalize across word position (5) and context (6), implying some degree of contextual invariance in brain representations of speech.

These results suggest a possible intermediate step that might be required in order for abstract categories to be learned: specifically, rather than learning categories directly from the acoustics, learners might first build representations of the speech signal that abstract away from contextual variability (48), and only later learn phonetic categories. Here, we simulate and extend results from Gwilliams et al. in order to examine whether such representational abstraction can occur in self-supervised models, and under what conditions. Given the difficulty that previous models have shown with learning abstract phonetic categories, evidence of learned invariance in the representations of any of the models is both surprising and informative.

We extended Gwilliams et al.’s analyses in three ways. First, we considered both cross-position and cross-context generalization. While Gwilliams et al. only studied the former, the latter was examined by Gennari et al. (6) using EEG recordings from 3-month-old infants. They found that decoders performed similarly when tested across phonetic context, although they used isolated syllables composed of six consonants and two vowels rather than continuous speech with a much richer repertoire of phonemes. Nevertheless, generalization across phonetic contexts is important to study due to the co-articulation effects mentioned above. For example, vowels are typically nasalized when followed by nasal consonants (43, 44), and plosives (stop consonants) can show spectral variation depending on the following vowel (45). In fact, behavioral studies addressing positional generalization have explicitly controlled for phonetic context (49). For completeness and comparability with Gwilliams et al., we studied both types of generalization using the same paradigm.

Second, we separated our analyses of vowels and consonants, which differ substantially in terms of distributional and co-articulatory properties. For example, in English the first phone of a word is more often a consonant, while vowels are more common at the second position. In terms of context, the co-articulatory influences also differ systematically: vowels are more often surrounded by consonants, and vice versa. Therefore, we separately analyze vowels (five monophthongs: Λ, ε, 3^˞^, I, i) and consonants (six plosives: p, b, t, d, k, g). For cross-position generalization, we consider five positions for vowels and plosives. For cross-context generalization, we consider six contexts for vowels, as defined by the manner of articulation of neighboring consonants, and four contexts for plosives, as defined by the front-backness of the preceding and the following vowel (see details in *Materials and Methods*).

Finally, we accounted for acoustic similarity when interpreting the decoding results. Acoustic signals can support at least some degree of generalization, since the acoustic realization of a phone often shares certain common acoustic properties across position and context. To evaluate the role of acoustic similarity, we ask how much generalization can be supported by acoustic features alone, and whether model representations can support more.

As adult neural data is only available for cross-position generalization (Figure 4(a)), we first examine how vowel and plosive decoders generalize across positions, simulating the brain data analysis by training decoders on p1 and testing on p2 and p3. Results are shown in Figure 4(b) for CPC-speech, CPC-audio, and acoustic features; results for Wav2Vec2 are in Figures S5 and S7.

**Fig 4.**
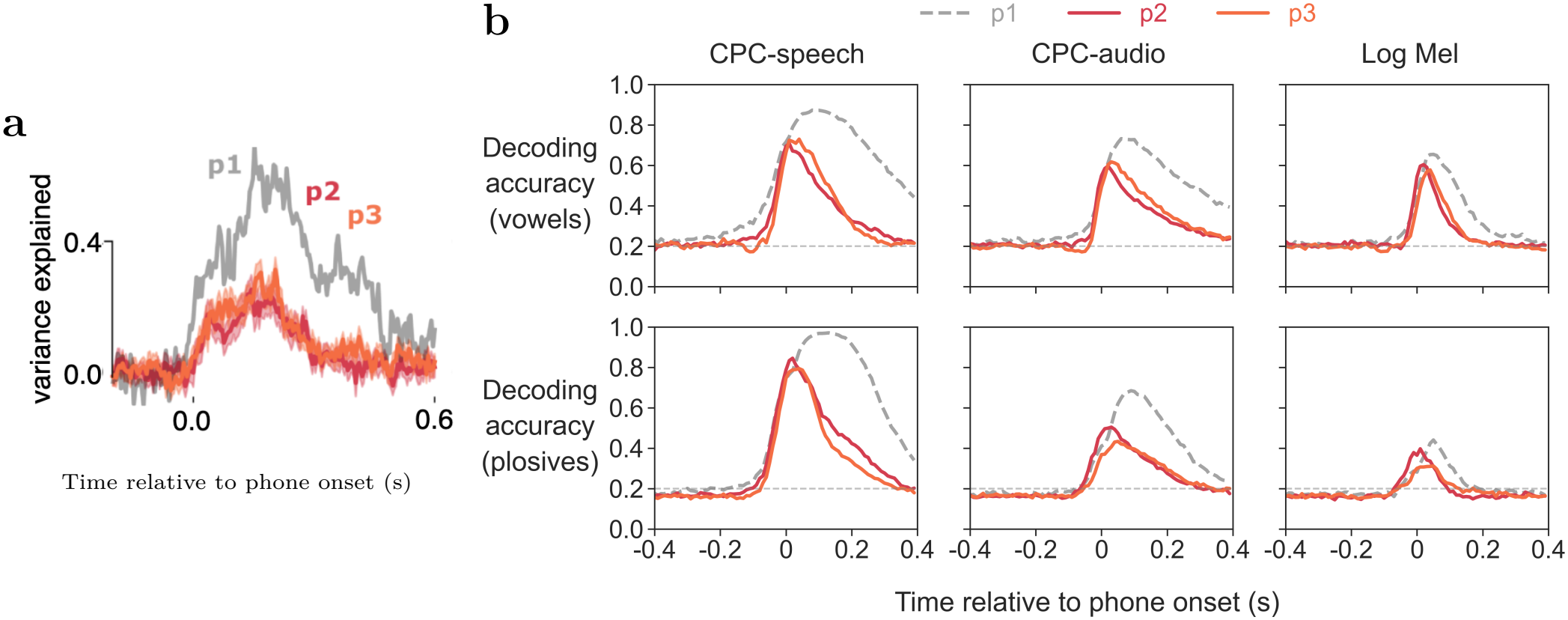
Cross-position generalization results, where decoders are trained on word-initial phones and tested on the second and third phones. (a) Gwilliams et al.’s results, decoding phonetic features of both vowels and consonants from MEG recordings. (b) Our results, decoding for five vowels (top row) and six plosives (bottom row) using representations from CPC-speech, CPC-audio, and acoustic features of log Mel spectrograms. The majority class baseline is shown as a dotted line.

Looking first at the results from CPC-speech, we see that for both vowels and plosives, the plots are qualitatively similar to the brain data: that is, the decoder trained at p1 generalizes relatively well to p2 and p3, but does show some drop in accuracy, particularly later during the decodable window. In other words, vowels and plosives at p1 remain decodable for much longer than those at p2 or p3. However, further analysis showed that for the phones in our analysis, and especially the vowels, their duration is longer wordinitially than elsewhere (average duration of the vowels we considered at p1: 110 ms, p2-p5: 83 ms; for the plosives p1: 95 ms, p2-p5: 80 ms). The longer window of decodability for p1 is therefore likely due to these differences in the acoustic duration at different positions. When we generalize beyond these vowels and consonants to look at all the phone classes, which together give a similar average duration for p1 and non-p1, the decodable window of p1 displayed a rightward shift but is not substantially longer (see Figure S3).

Due to these issues with interpreting the width of the decodable window, in the remaining analyses we consider only the maximum accuracy of the decoders in each position or context. We quantify generalization by comparing the maximum accuracies (relative to the baseline obtained by predicting the majority class) across different conditions. More specifically, given a particular model (or acoustic features), *gen strength(A → B)* shows how well a decoder trained on that model’s representations from condition A (a particular position or context) performs on an unseen condition B. It is computed as (*acc*(*B*) *− c*)*/*(*acc*(*A*) *− c*), where *acc*(*X*) is the peak accuracy under condition *X*, and *c* is the baseline (always 0.2 for vowels and 1/6 for plosives, since we balanced the classes for the analyses in this section; see *Materials and Methods*). A score of 100% indicates complete generalization, whereas a score of 0% means the decoder only performs at the baseline level under condition B.^‡^ Using this measure, each panel of Figure 4(b) will be reduced to two numbers: *gen strength(p*_1_ *→ p*_2_*) and gen strength(p*_1_ *→ p*_3_).

We can now move beyond our qualitative replication of the brain data, which showed that models partially generalize across positions, to ask whether they do so to a greater extent than would be expected due to acoustic similarity. We do so by comparing the generalization strength of each model on each condition to that of the acoustic features. For example, looking at vowels, *gen strength*(*p*_1_ *→ p*_2_) using CPC-speech is (0.71 *−* 0.2)*/*(0.87 *−* 0.2) = 0.75, and using log Mel features it is (0.6 *−* 0.2)*/*(0.65 *−* 0.2) = 0.87. For vowels with this particular pair of training–testing positions, acoustic features actually support *better* generalization than model representations. This example underscores our argument that analyzing generalization in model (or brain) representations alone is not enough to support claims of invariance. Still, this is only one of many possible training–testing pairs, so to assess generalization more broadly, we need to analyze all possible pairs.

Figure 5 shows the results of this analysis for both vowels and consonants. We first consider the results for cross-position generalization in vowels (upper left). The plot shows *gen_strength*(*p*_*i*_ *→ p*_*j*_) of vowel decoders for all combinations of *i* and *j* (*i, j ∈* [1, 5], *i* = *j*). This results in 20 markers for each model, where the *y* value of each marker represents *gen_strength* of model representations, and the *x* value is *gen_strength* of acoustic features under the same training and test positions. If the *gen strength* of a model is the same as that of acoustic features, the corresponding marker would fall on the diagonal. In the case of vowels across positions, most markers fall near or below the dashed diagonal line. This suggests that generalization based on model representations is similar to, if not slightly weaker than that of acoustic features. The upper right figure shows similar results for generalization of vowel decoders across contexts. Overall, model representations and acoustic features both exhibited a medium to high degree of generalization across different positions and contexts. This means that vowels are largely and inherently position- and context-invariant, and there is little evidence that models are learning more abstract representations.

**Fig 5.**
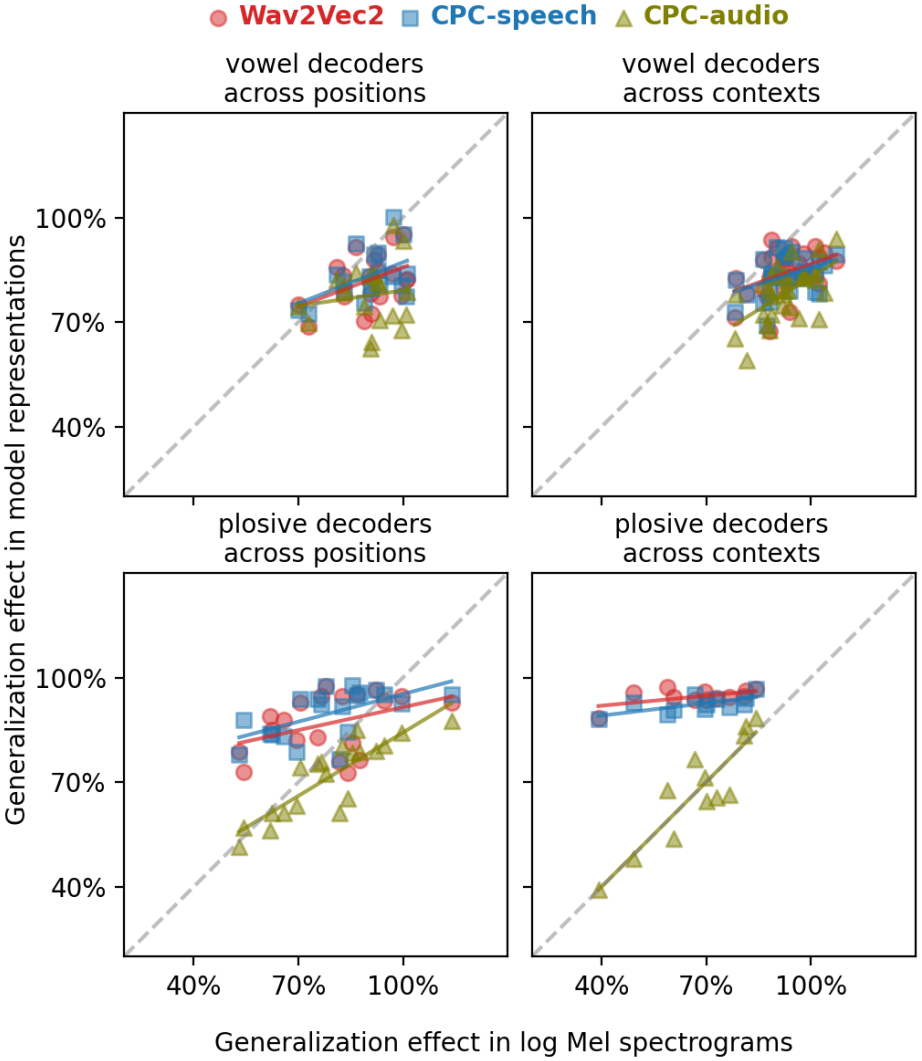
Generalization strengths of model representations (each model designated with a different color) against those of acoustic features. Each marker represents a single combination of training and testing conditions (for example, training on position 1 and testing on position 2), indicating *gen strength* of model representations (*y* value of each marker) and *gen strength* of acoustic features (corresponding *x* value). The decoding accuracy curves for all pairs of training and test conditions, from which these results were computed, can be found in Figures S5–S8.

However, a very different picture emerges for plosives, as we can see from the two bottom panels. Plosives are more variable acoustically: the range of *gen_strength* based on acoustic features is much greater (x-axis values vary between 50% and 110% across positions, 40% and 100% across contexts), and this variation is mirrored in CPC-audio. Yet the two speech models generalize much better: Wav2Vec2 and CPC-speech maintain high *gen_strength* across positions and near-perfect generalization across contexts.

Through statistical analyses (see *Materials and Methods*), we further examined the advantage of speech models over CPC-audio. On average, the speech models support better generalization than CPC-audio, with a stronger effect for plosives than for vowels (see Figure S10 for details). The strongest support for abstraction would come from cases where acoustic generalization is weak. We find that for plosives, but not for vowels, the difference in generalization between speech and audio models is highest in this circumstance, indicating that the models are even abstracting across positions and contexts where plosives are acoustically most dissimilar.

To confirm that the generalization advantage of the speech models is due to training on speech, rather than their larger training corpora relative to CPC-audio, we also tested a second CPC-speech model with an identical architecture to CPC-audio but trained on about one-fifth as much data. This model showed a similar generalization advantage over CPC-audio (Figure S9). This suggests self-supervised learning from non-speech audio is insufficient to overcome the variability in the acoustics of plosives, but models exposed to speech can learn to abstract away from acoustic variations, creating a more stable, invariant representation.

Taken together, our results show that self-supervised learning from speech can give rise to representations that abstract over positional and contextual variability, at least for some sound classes. These representations retain information about preceding and (for Wav2Vec2) following phones, while also disentangling each phone from its neighbors, paving the way for fully abstract, discrete categories. Such representations were not found in the model trained on non-speech audio, where generalization largely reflected acoustic similarity. Methodologically, our results underscored the importance of accounting for acoustic similarity when analyzing phonetic representations in brains (50), and of separately analyzing different phonetic categories. Further analyses of neural signals are required to determine the extent of invariance in speech representations of human listeners.

## Discussion

Our results shed light on how self-supervised learning and exposure to speech input shape the properties found in humans’ neural representations of speech. We found that self-supervised learning models exhibit the same kind of extended decodability and dynamically evolving encoding patterns that have been found in brain representations of speech. We showed that these properties, which have been hypothesized to support the integration of speech sounds over time, can arise from self-supervised learning over general auditory input, even without exposure to speech. Beyond temporal dynamics, we developed a measure of context-invariance that takes into account acoustic similarity, and showed that only the models trained on speech data were able to develop context-invariant representations. Such abstraction does not arise from self-supervised learning alone, nor simply from encoding context, as the audio model does both, yet still cannot support cross-context generalization beyond what is afforded by the acoustic signals.

We showed that dynamic neural codes can arise without requiring knowledge of higher-level units or external feedback beyond the speech signals. Our finding does not necessarily mean the models have not induced any higher-level structures — the task of predicting upcoming acoustics necessarily encourages the model to capture temporally-extended regularities or structures. However, any such structures would be non-linguistic for the audio model and need not correspond to accurate word-level units for the speech models. This finding is potentially relevant for explaining how similar temporal generalization patterns could arise in the EEG data of 3-month-old infants before they know words (6).

The functions served by dynamic neural codes may also be domain-general. Apart from the speech domain, they have also been found in the brain signals of people listening to auditory tones (51) or watching visual stimuli presented in fast sequences (52). Together, these results support the idea that dynamic neural codes reflect general principles of sensory processes, many of which involve order-sensitive integration of sequential input. On the other hand, it has also been found that language processing invokes not only dynamic neural codes but a hierarchy of such encoding (53). A potential future application of our modeling framework could be to examine which parts of the hierarchical encoding can emerge from self-supervised learning, and which require domain specificity in the input data or the learning algorithm.

In contrast to the domain-generality of temporal dynamics, we attributed the abstraction over context to self-supervised learning over speech input specifically. While linguists have long posited the presence of abstract phonological categories (43, 54), co-articulation effects cause the acoustic realization of such categories to be highly variable (55), and computational models have struggled to recover abstract categories from realistic, continuous raw speech (46, 47). Our results suggest self-supervised learning from speech can link acoustically diverse instances of the same phoneme (i.e. allophones) to the same category, at least for certain phonemes and contexts. How this invariance is achieved, and whether the model approximates previous proposals regarding phonological learning, e.g. distributional learning or proto-lexicon building (56, 57), is an interesting question for future research. Future work could also examine whether abstraction is induced for other well-studied allophones, such as place-assimilated nasals (58).

It remains uncertain whether this kind of context-invariance is also present in brain representations, and if so, how the brain data would compare with the speech and the audio models in this respect. Gwilliams et al. did not examine generalization across phonetic contexts. They did find partial generalization across positions; however, since they did not compare the degree of generalization achieved by the brain representations to an acoustic feature baseline, we don’t know whether the brain representations are achieving higher invariance than would be expected based on acoustic similarity alone. Moreover, their analyses aggregated across all phonetic features, whereas our modeling results indicate that contextual invariance may differ across phonetic categories. To fully compare our analyses to brain representations, those generalization results would need to be broken down by phonetic features or categories, and generalization would need to be measured across phonetic contexts in addition to word position.

Interestingly, the pattern of context dependence we found in the non-speech audio model resembles what Gennari et al. found in the EEG data of infants (6). Although their study used isolated syllables rather than continuous speech, they also found greater context sensitivity for consonant place of articulation than for vowel identity, as we observed in the audio model. The greater context variability in consonants is also consistent with the well-established finding that phonetic learning of vowels precedes that of consonants during language acquisition (59–61). While additional simulations are needed to determine whether the audio model can simulate all of Gennari et al.’s findings, this parallel highlights the value of the audio model as a tool for exploring how much of phonetic discriminability or which stage of phonetic learning in infants can be explained by general auditory mechanisms.

Although this work is primarily aimed at cognitive scientists, it is also relevant for speech technology researchers, since similar models are used as the basis of many current systems. Within the speech technology community, analyses of models’ phonetic representations have mainly focused on comparing phone decodability across different models or model layers (12, 62–68). While there is some work on *how* such information is encoded (15, 69–73), we know of no analyses examining the temporal dynamics of speech representations, or of any that investigate context-invariance by testing decoders for generalization to unseen contexts. These questions and methods, inspired by work in cognitive neuroscience and demonstrated on SSL models by this study, could prove fruitful for understanding and improving these models.

To conclude, our work fits into the larger landscape of research using computational models to answer *why* the brain works the way it does (74). There has been a surge of studies comparing deep neural network models and brains on speech and language processing, which have primarily focused on global correlations between brain signals and model representations (“brain scores”) (21–28). Such analyses have been criticized for leaving substantial ambiguity regarding what drives the correlation (29, 30). We focused instead on directly simulating the neural signatures that are hypothesized to support a particular cognitive function (speech recognition), which allowed us to tease apart which aspects of the representations could be attributed to different factors—self-supervised learning, high dimensional representations, acoustic similarity, or domain-specific input—rather than simply comparing overall correlations between models. As a powerful and complementary tool for neuroscience, our modeling approach paves the way for a more concrete understanding of the neural mechanisms underlying speech processing.

## Materials and Methods

### Models

In terms of model architecture, both CPC and Wav2Vec2 are composed of an encoder network and a context network. The encoder network takes a raw audio waveform as input, and maps it to a series of vector representations. These representations are then contextualized by being passed through the context network. In training, the contextualized representations are used to predict the encoder output by optimizing a contrastive loss. A key difference between CPC and Wav2Vec2, though, lies in their prediction targets: Wav2Vec2 discretizes the continuous encoder outputs through vector quantization before using them as targets, whereas CPC directly predicts the continuous latent features. In addition, Wav2Vec2 uses a diversity loss to encourage the model to use all discrete targets equally.

In both CPC and Wav2Vec2, the encoder is a convolutional neural network, although with different number of layers (both CPC-speech and CPC-audio: 5, Wav2Vec2: 7) and dimensionality (CPC-speech: 512, CPC-audio: 256, Wav2Vec2: 768). These representations have a temporal resolution much lower than the raw audio signals (10 ms for CPC and 20 ms for Wav2Vec2), but comparable to that of log Mel spectrograms (10 ms). The “receptive field” of each representation, which is the span of the input waveform that affects a single representation, is 29ms for CPC-speech and CPC-audio and 25ms for Wav2Vec2, due to different strides (CPC: 5,4,2,2,2, Wav2Vec2: 5,2,2,2,2,2,2) and kernel widths (CPC: 10,8,4,4,4, Wav2Vec2: 10,3,3,3,3,2,2). For the context network, which is jointly trained with the encoder, CPC uses LSTM layers (CPC-speech: 4 layers, CPC-audio: 2 layers) whereas Wav2Vec2 applies 12 transformer layers.

The CPC-speech model was implemented and trained by (31) on 6000 hours of audiobooks from the “clean-light” subset of the Librilight corpus (75). We used representations from the output of the second layer, which gave the highest decoding accuracy (31). The Wav2Vec2 model is the “base” model in (34), which was trained on 960 hours of audiobooks from the “LS-960” subset of the Librispeech corpus (76). We used representations from the output of the ninth layer, the layer that encodes most phonetic information (68). The CPC-audio model was obtained from (32). The training set includes 78 hours of field recordings of animals from the Animal Sound Archive (77), and 422 hours of animal sounds or everyday environmental sounds from Audioset (78). We used representations from the output of the second layer.

For Figure S9, we also used a smaller CPC-speech model, provided by the same group that trained the aforementioned CPC-speech (31). It has the same architecture as CPC-audio, with identical dimensionality, number of layers, and other hyper-parameters. It was trained on 100 hours of speech from the “train-clean-100” subset of Librispeech.

The log Mel spectrograms were extracted with torchaudio.compliance.kaldi.fbank with the default parameters. The smoothed log Mel features were computed using exponential smoothing, with the smoothing factor set to 0.3.

### Data

In the first and second simulations, for training and testing the decoders, we used the speech audio from the *dev-clean* subset of Librispeech (76). It contains audiobooks read by 21 male and 19 female speakers, with 8 minutes of speech from each speakers. For each chapter read by each speaker, we split the utterances in half to make up the training and the test utterance sets. This results in 1329 utterances (~ 154 minutes) in the training set and 1374 utterances (~ 168 minutes) in the test set.

In the third simulation, we used the *train-clean-360* subset of Librispeech, which is substantially larger than *dev-clean* (360h of speech). We used this larger subset in order to obtain enough samples of each phone in each position/context to robustly train and test the decoders. We note that this subset overlaps with the training data of Wav2Vec2 (though not the CPC models); however, the overlap is unlikely to affect our conclusions, since we found qualitatively similar (but much noisier) results using *dev-clean*, which was not part of the Wav2Vec2 training data.

The phone labels and boundaries required for our anlayses were obtained by performing forced alignment with an acoustic model created according to the official Kaldi recipe for LibriSpeech data (https://github.com/kaldi-asr/kaldi/blob/master/egs/librispeech/s5/run.sh). There are 39 phoneme categories, including

- 15 vowels: α, ae, ʌ, ɔ, aʊ, aɪ, ɛ, 3^˞^, eI, I, i, oʊ, ɔ?, ʊ, u
- 24 consonants: m, n, ŋ, ɹ, l, j, w, p, b, t, d, k, g, d_3_, h, f, v, s, z, θ ð, ʃ, tʃ, 3

Further filtering of the data was done for each simulation, as detailed in the respective sections below.

### Time-resolved decoding

Time-resolved decoding involves grouping representations according to their relative latency to a reference time point, which is the phone onset in our simulations, and training a separate decoder for each group. We trained multinomial logistic regression models as decoders, using the implementation from the scikit-learn. The SlidingEstimator function from the MNE package (79) allows us to perform time-resolved decoding in an efficient manner. The decoders are optimized with the LBFGS solver till convergence or reaching 100 iterations, and evaluated using accuracy as the metric.

### Decodable window

In this and the rest of our simulations, we excluded words with only one phone, resulting in 24940 phone samples in the training set and 27246 phone samples in the test set. For this simulation, we considered a time window of 1200 ms centered on phone onset. The window is padded with all-zero vectors as needed when a phone occurs in the beginning or the end of the audio file. 2% of the phone samples required padding to the left, and 4% to the right. We train 120 decoders for CPC-speech and CPC-audio respectively (60 for Wav2Vec2 which has half the sample rate), one on each relative timestep, and evaluate them on the same timesteps in testing.

### Temporal generalization

We considered a time window of 600 ms, which starts 200 ms before phone onset and ends at 400 ms after onset. This means training 60 decoders for CPC-speech and CPC-audio respectively, and 30 for Wav2Vec2. The temporal generalization test differs from the previous simulation in that each decoder is tested not just on the group with the same latency as it has seen in training, but on all of the groups. For example, the decoder trained for representations occurring at 100 ms before onset will be tested on each of 60 groups of representations, resulting in 60 accuracy values. Applying the same tests to all the 60 decoders results in a 60 *×* 60 matrix.

As the temporal generalization results are aggregated separately for each phone position, we obtain eight matrices, which are superimposed and presented in the same figure through appropriate arrangement of each matrix. To align the matrix corresponding to each phone position according to the average timeline of phones in a word, we adjusted their relative positions as follows: matrices for p2–p4 were incrementally shifted to the right by the average duration of the preceding phone, while matrices for p-1–p-4 were shifted to the left by the average duration of the following phone.

### Generalizing decoders across position or context

For vowels, we considered the five most common monophthongs: Λ, ε, 3^˞^, I, i. The positions considered are from the first to the fifth phone in a word, and the contexts are: plosive__fricative (i.e. vowels preceded by a plosive and followed by a fricative), plosive__plosive, plosive__nasal, fricative__fricative, fricativeplosive, fricative__nasal.

For plosives, we considered p, b, t, d, k, g. The positions considered are from the first to the fifth, and the four contexts are defined as: front__front, front__back, back__front, back__back, where front vowels include i, I, ε, and back vowels include α, Λ, 0Λ8α, u, ⊃.

For each vowel (or plosive) occurring in each position (or context) that we consider, we sampled 500 samples from the training utterances, and used them for training the decoders. Similarly, 500 samples are sampled from each category for testing generalization of the decoders. This ensures the decoders are trained and tested with balanced target classes, as we noticed non-uniform class distribution can lead to overestimation of generalization strengths.

### Statistical analysis

For each of Wav2Vec2, CPC-speech, and CPC-audio, we fitted an ordinary least-squares regression using the *statsmodels* package in python. The dependent variable was the difference in generalization strength between the model of interest and CPC-audio, and the predictor was mean-centered generalization strengths of acoustic features. The values for the intercept and the slope are reported in Figure S10.

### Preprocessing

Gwilliams et al. pointed out that phonetic information is partly confounded with low-level acoustic properties such as amplitude and pitch, and therefore they preprocessed the neural recordings using a linear model to regress out these two factors. For model representations, we tried performing preprocessing by training two ridge regression models to predict amplitude and pitch values from each representation, and regressing out the variance in the direction given by the regressors’ coefficients. We found that this operation made negligible difference to the results of the simulations.

## ACKNOWLEDGMENTS

This work was supported in part by the UKRI Centre for Doctoral Training in Natural Language Processing, funded by the UKRI (grant EP/S022481/1) and the University of Edinburgh. We thank Rebecca Bull for running preliminary experiments as part of our invariance analysis, and Rebecca Bull and Stefano Coretta for their help in designing the statistical model for that analysis.

## Supporting Information

### Decodable window: control for stimulus dependency

There are three different types of information that could help a decoder predict the phone label. First, *acoustic information* about the phone, including from coarticulation, will primarily be available during a short window around the phone itself. Second, speech contains strong *stimulus dependencies* because some sequences of phones are common and others are rare or impossible: for example, if the acoustic information available at –150ms is most consistent with the sequence [st], then the phone starting at 0ms is likely to be a vowel or [r], and a decoder could predict that even before any acoustic cues to that phone are available. This is an example of *prediction*, but stimulus dependencies can also be used for *postdiction* (or backward prediction). That is, even if representations do not maintain information about the previous phone, a decoder could use stimulus dependencies to predict that phone based on the representation of a later phone. Therefore, although stimulus dependencies on average permit a wider window of decoding than coarticulation, we expect both of them to have a roughly symmetrical window. We confirmed this by introducing a control, *posteriorgrams*.

A posteriorgram is the categorical distribution over all the phoneme categories, which represents a model’s belief about the current phone. To obtain posteriorgrams, we trained a single decoder to predict the current phoneme category based on one representation, which might be sampled from any point during a phone. Once trained, the decoder was applied to obtain a posteriorgram for each frame, which has the same dimension as the number of phoneme categories (39). We then applied the decodable window analyses to the posteriorgram features.

**Fig. S1.**
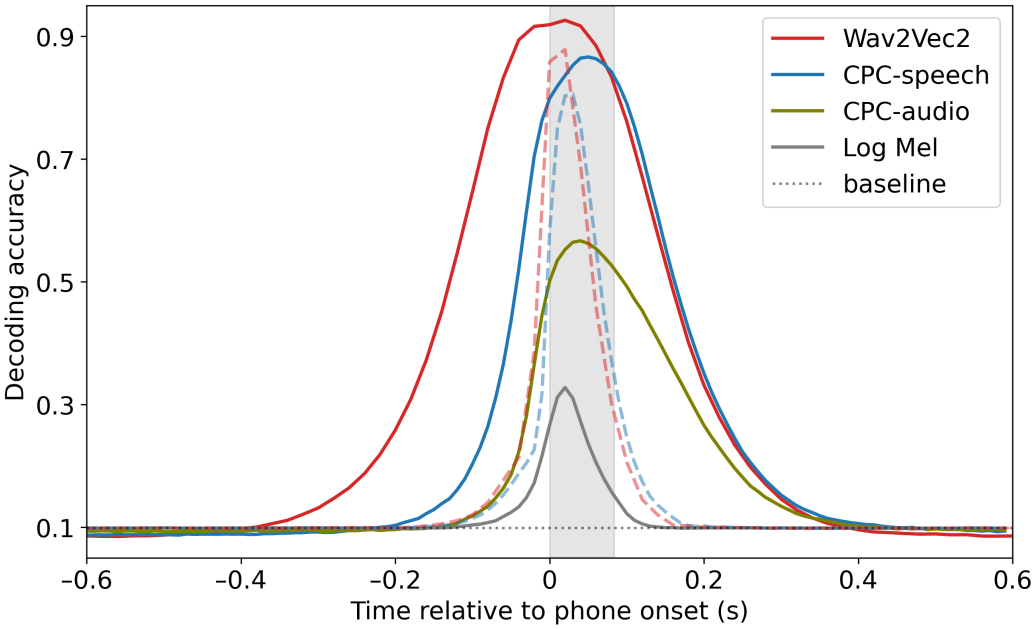
Decoding accuracy obtained with posteriorgrams based on model representations was plotted as the dashed curves.

For both CPC-speech and Wav2Vec2, we found that posteriorgrams support decoding for up to 200ms before and after phone onset (roughly 2-3 phones). Both curves are much more symmetrical around the phone onset than the solid curves, obtained with the full representations. The decodable windows are also substantially shorter than the range obtained with the original representations, which extends more than 200ms beyond that. These results suggest that most of the post-offset decodability we observe is not simply due to stimulus dependencies, but reflects explicit maintenance of prior phone information in the representations.

### Decodable window: control for representation dimensionality

One disadvantage of the log Mel spectrograms as compared to the model representations is the lower dimensionality. We thus reduced the model representations to the same low dimensionality using PCA and then evaluated the decodable window.

**Fig. S2.**
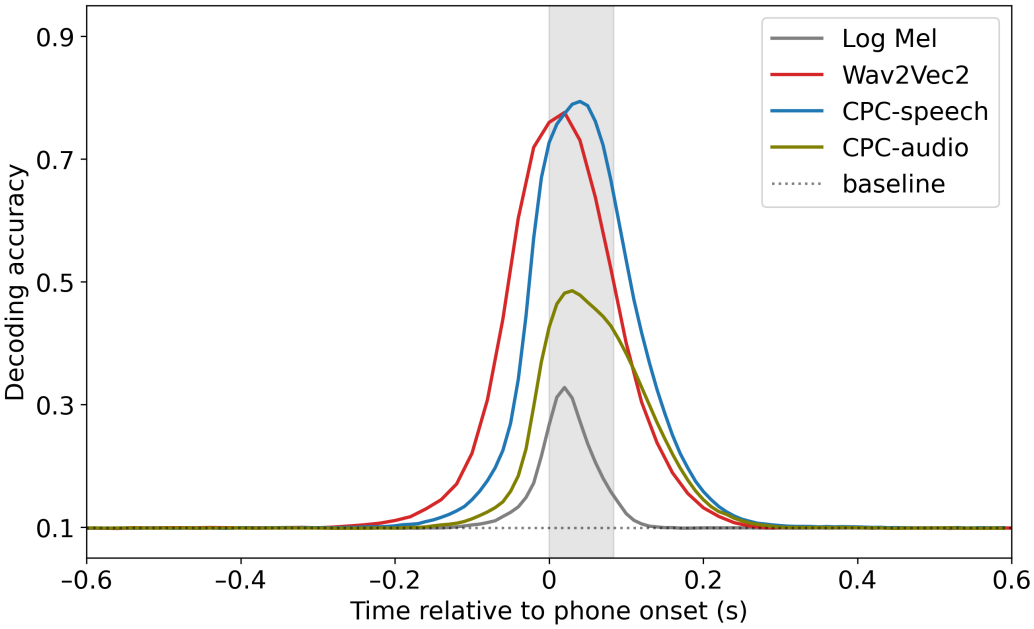
Decodable window of reduced-dimensionality representations.

Even with only 40 dimensions, model representations support wider decodable windows and higher decoding accuracy than in the acoustic features, though in this case, the difference between models and acoustics is much bigger in maintenance effect than in predictive effect. This suggests the higher principal dimensions capture more information about previous than upcoming phones.

### Decodable window: comparison between word-initial and non-initial phones

**Fig. S3.**
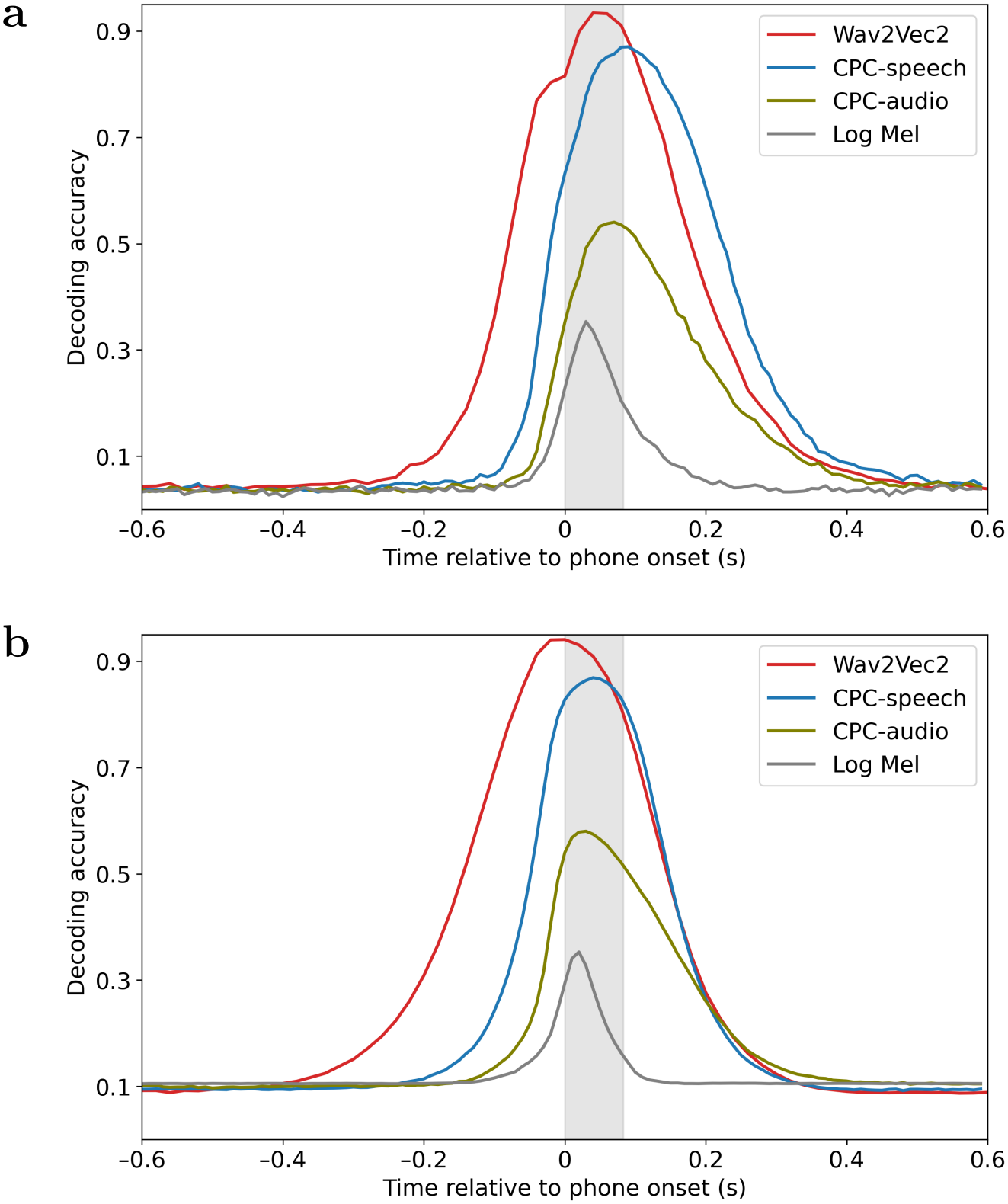
Decodable window for (a) word-initial phones (b) non-initial phones.

### Temporal generalization controls

**Fig. S4.**
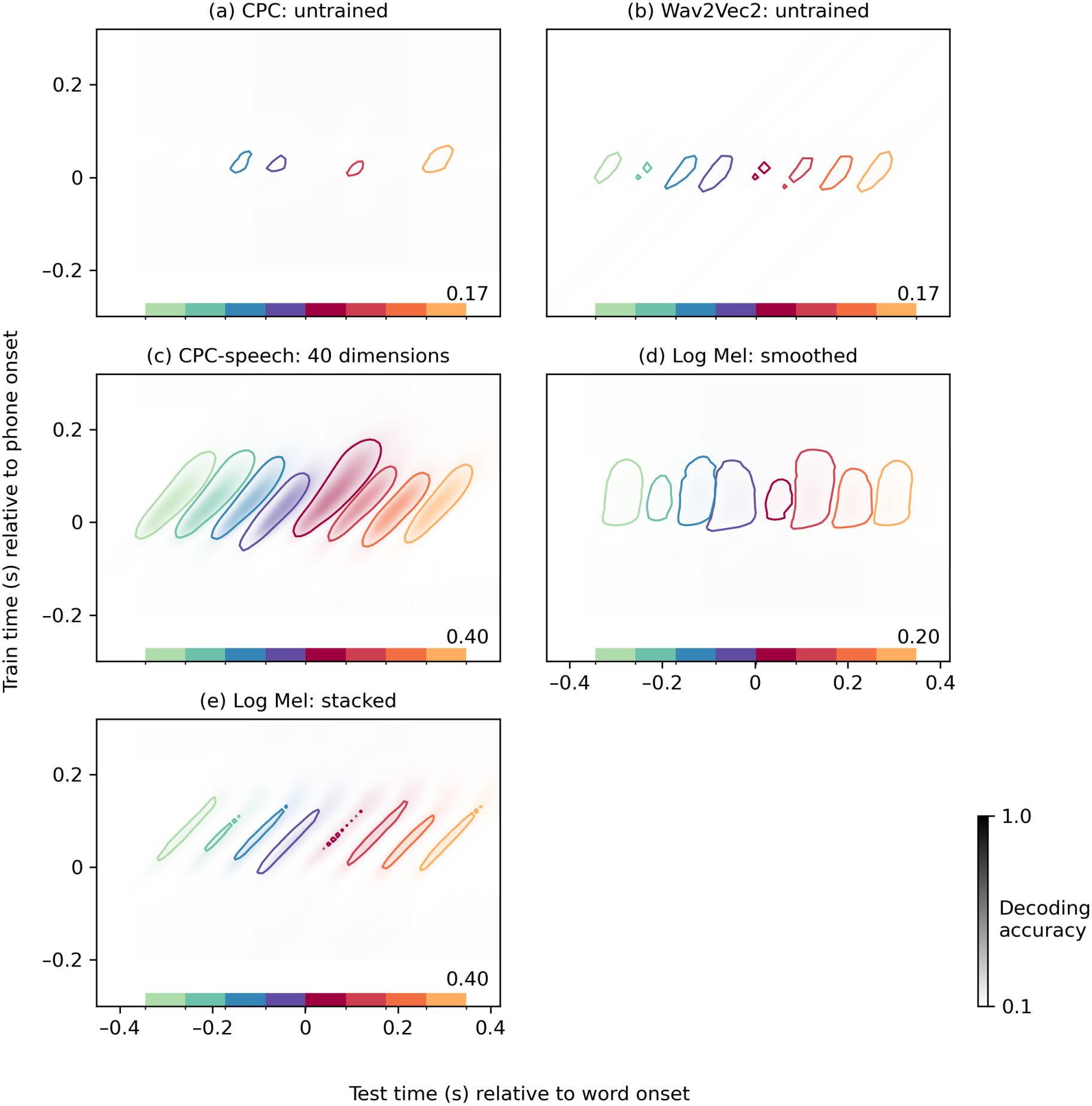
Temporal generalization results for various control representations. (a) Representations extracted from a randomly initialized, untrained CPC model with the same architecture and parameter size as CPC-speech. The contours were plotted at 0.17 accuracy. (b) Representations extracted from a randomly initialized, untrained Wav2Vec2 model (contours 0.17 accuracy). (c) Representations were extracted from CPC-speech and then reduced to 40 dimensions using PCA (contours at 0.4 accuracy, as in the plots in the main text). (d) Smoothed log Mel spectrogram representations (also 40 dimensions), using exponential smoothing with the smoothing factor set at 0.3. (e) Stacked log Mel spectrogram features, obtained from concatenating 13 single-frame features (13 *×* 40 = 520 dimensions, contours at 0.4 accuracy).

### Cross-position and context generalization

**Fig. S5.**
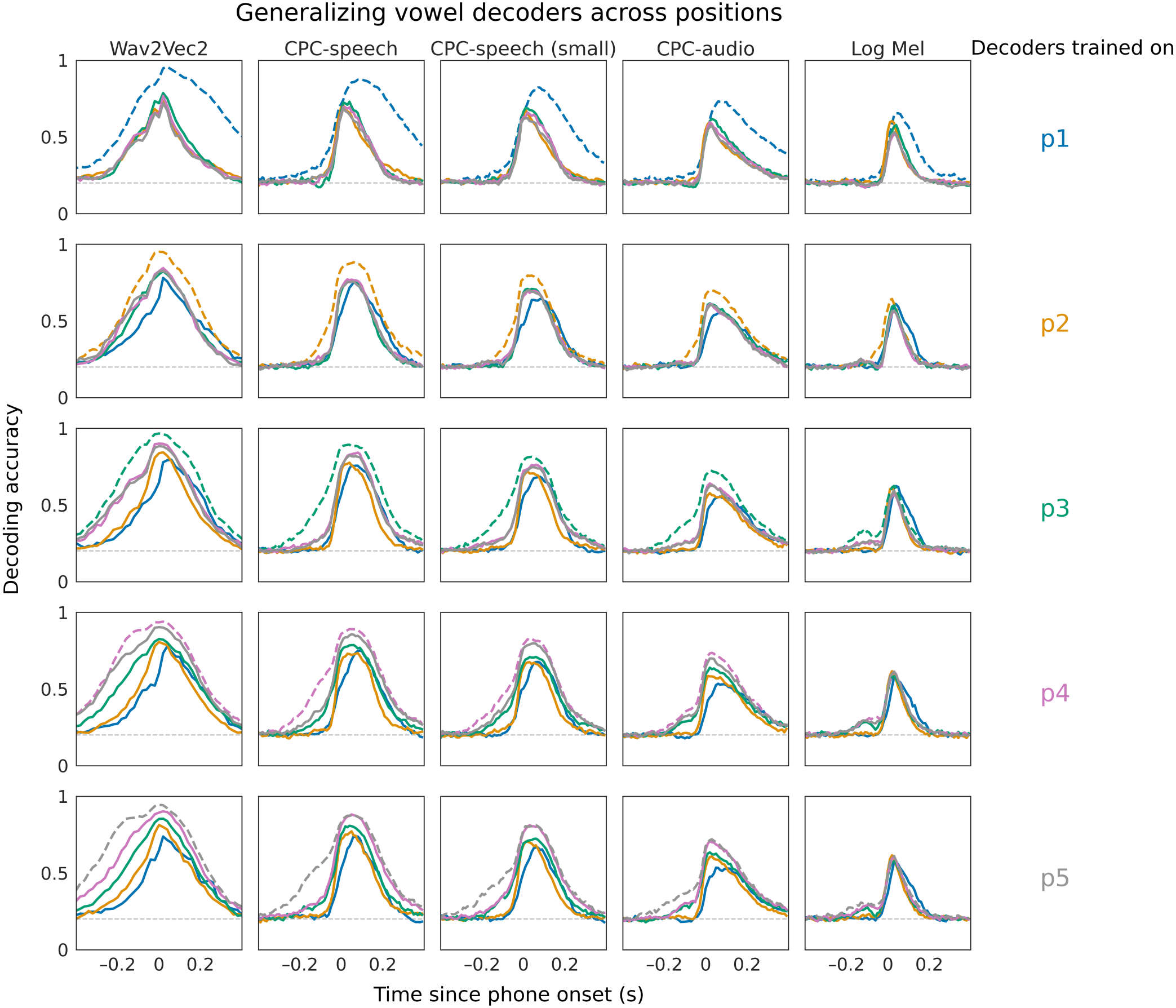
Generalizing decoders of five vowels (Λ, ε, 3, I, i) across five positions in the word. Each column of plots shows results from one model, and each row shows results from the same training position (labelled on the right). Each plot contains one dashed curve, which shows the accuracy when the decoders are tested on the same phone position it was trained on (with a different set of samples), and four solid curves, each showing the accuracy when the decoders are tested on samples occurring in an unseen position. The colors denoting each position are shown on the right.

**Fig. S6.**
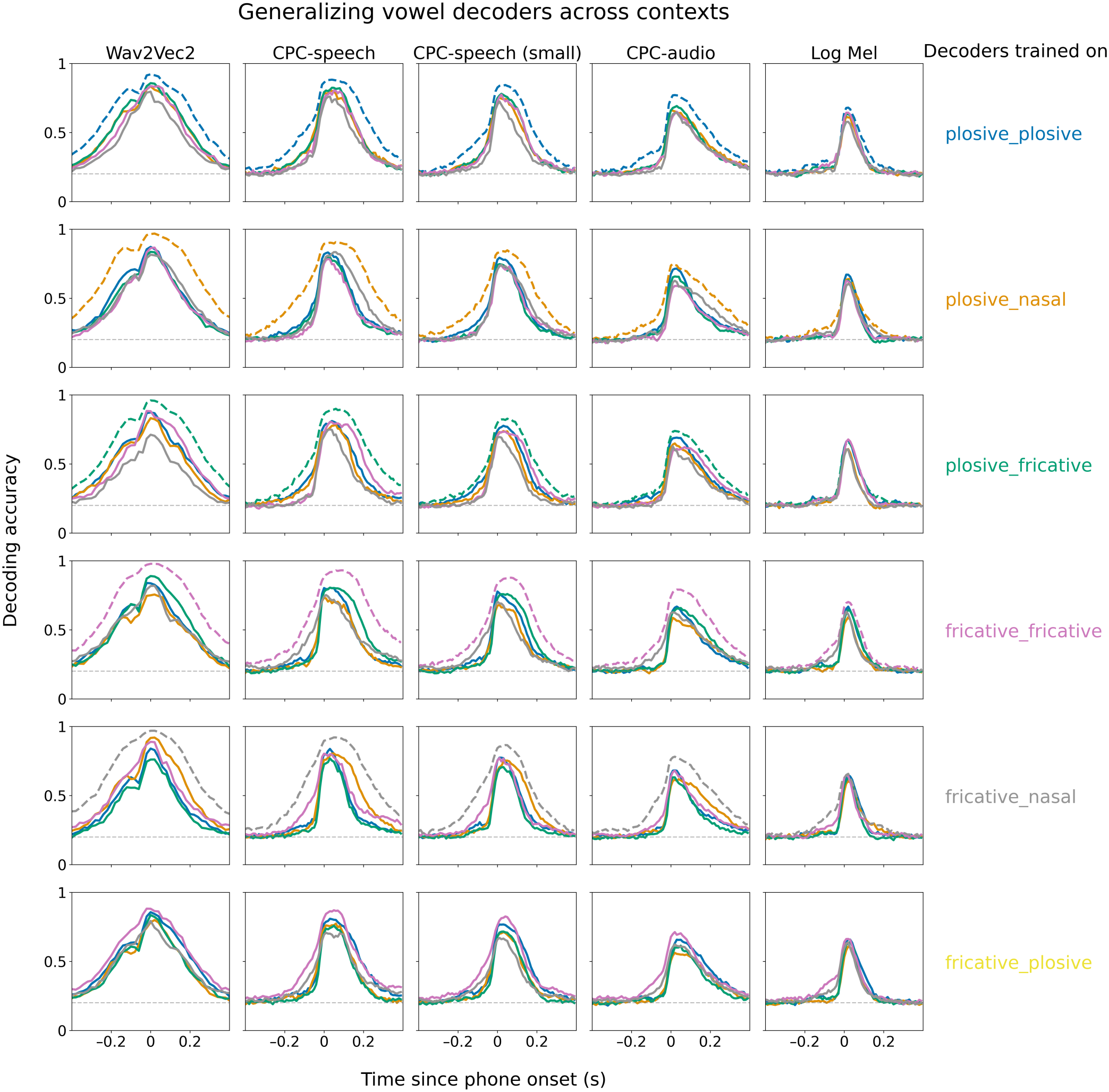
Generalizing decoders of five vowels (Λ, ε, 3, I, i) across six phonetic contexts as defined by the manner of articulation of the previous and subsequent phone. Each column of plots shows results from one model, and each row shows results from the same training context (labelled on the right). Each plot contains one dashed curve, which shows the accuracy when the decoders are tested on the same context it was trained on (with a different set of samples), and five solid curves, each showing the accuracy when the decoders are tested on samples occurring in an unseen context. The colors denoting each context are shown on the right.

**Fig. S7.**
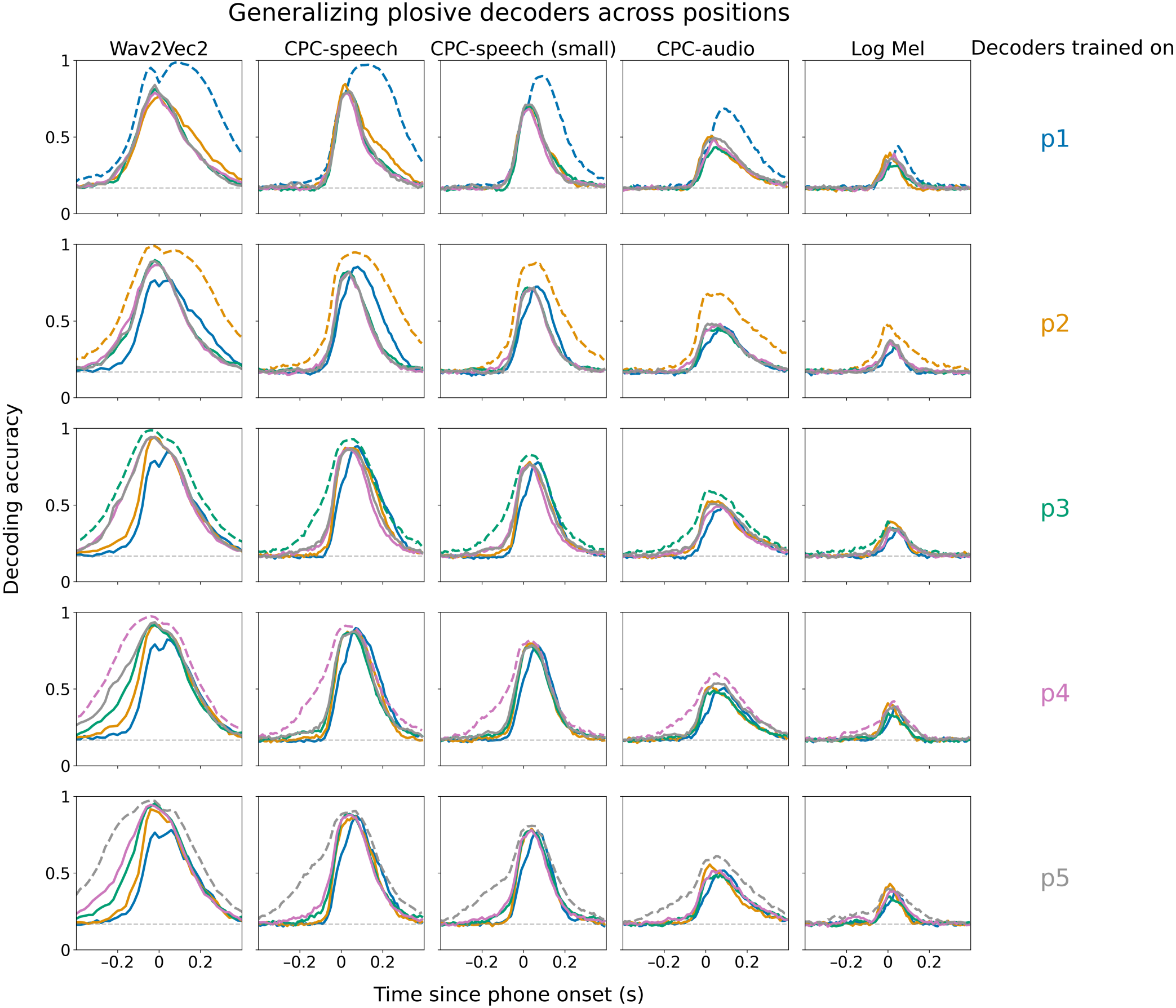
Generalizing decoders of six plosives (p, b, t, d, k, g) across five positions in the word. Each column of plots shows results from one model, and each row shows results from the same training position (labelled on the right). Each plot contains one dashed curve, which shows the accuracy when the decoders are tested on the same position it was trained on (with a different set of samples), and four solid curves, each showing the accuracy when the decoders are tested on samples occurring in an unseen position. The colors denoting each position are shown on the right.

**Fig. S8.**
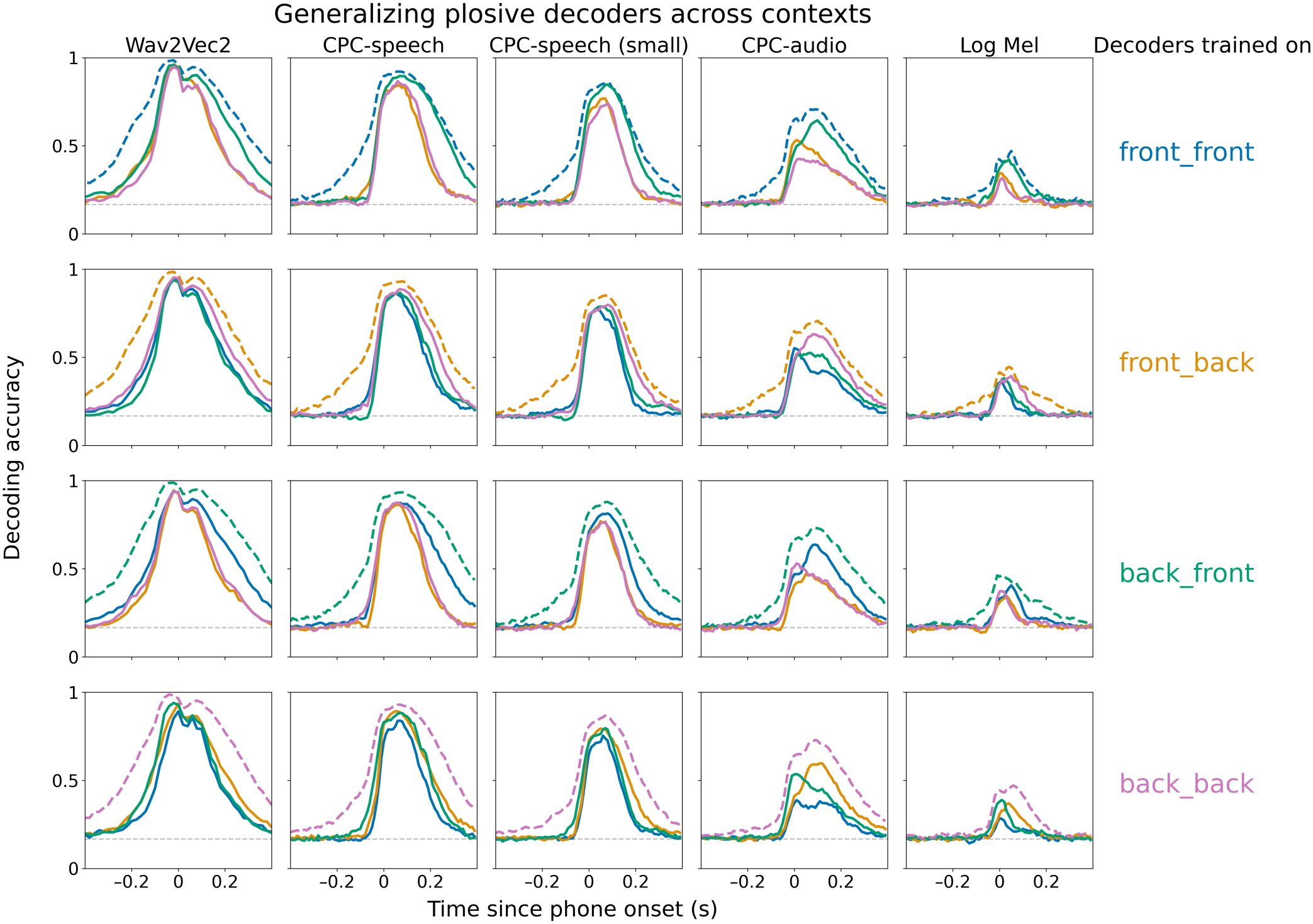
Generalizing decoders of six plosives (p, b, t, d, k, g) across three contexts, as defined by the front-backness of the preceding and the following vowel. Each column of plots shows results from one model, and each row shows results from the same training context (labelled on the right). Each plot contains one dashed curve, which shows the accuracy when the decoders are tested on the same context it was trained on (with a different set of samples), and three solid curves, each showing the accuracy when the decoders are tested on samples occurring in an unseen context. The colors denoting each context are shown on the right.

**Fig. S9.**
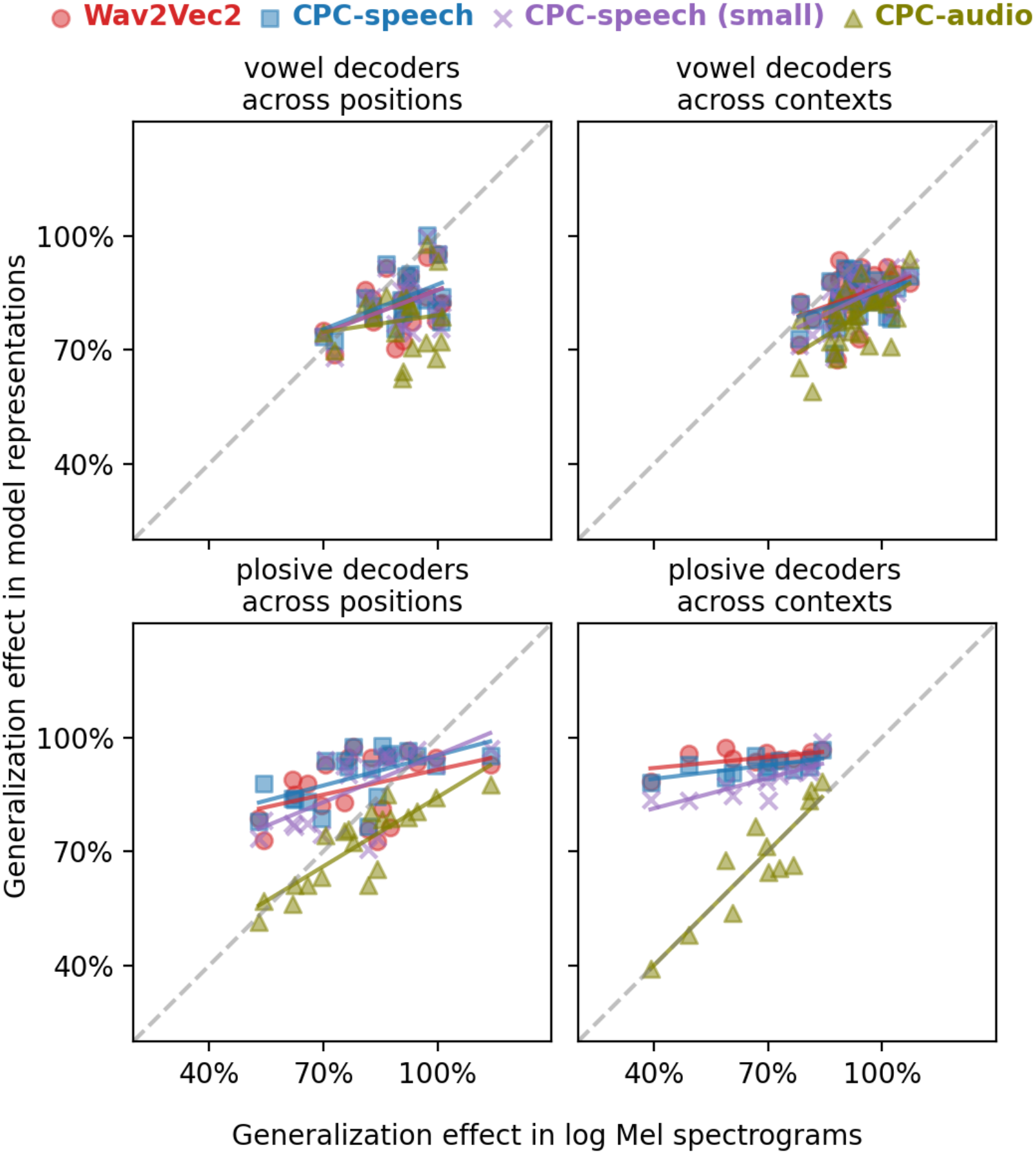
Generalization strengths of model representations (plotted along y-axis) and of acoustic features (x-axis). Compared to Fig. 5 in the main paper, this version includes the CPC-small model, which patterns like the other two speech models, i.e. it exhibited a higher degree of invariance for plosives than CPC-audio despite having an identical architecture and trained on a fifth of the data.

**Fig. S10.**
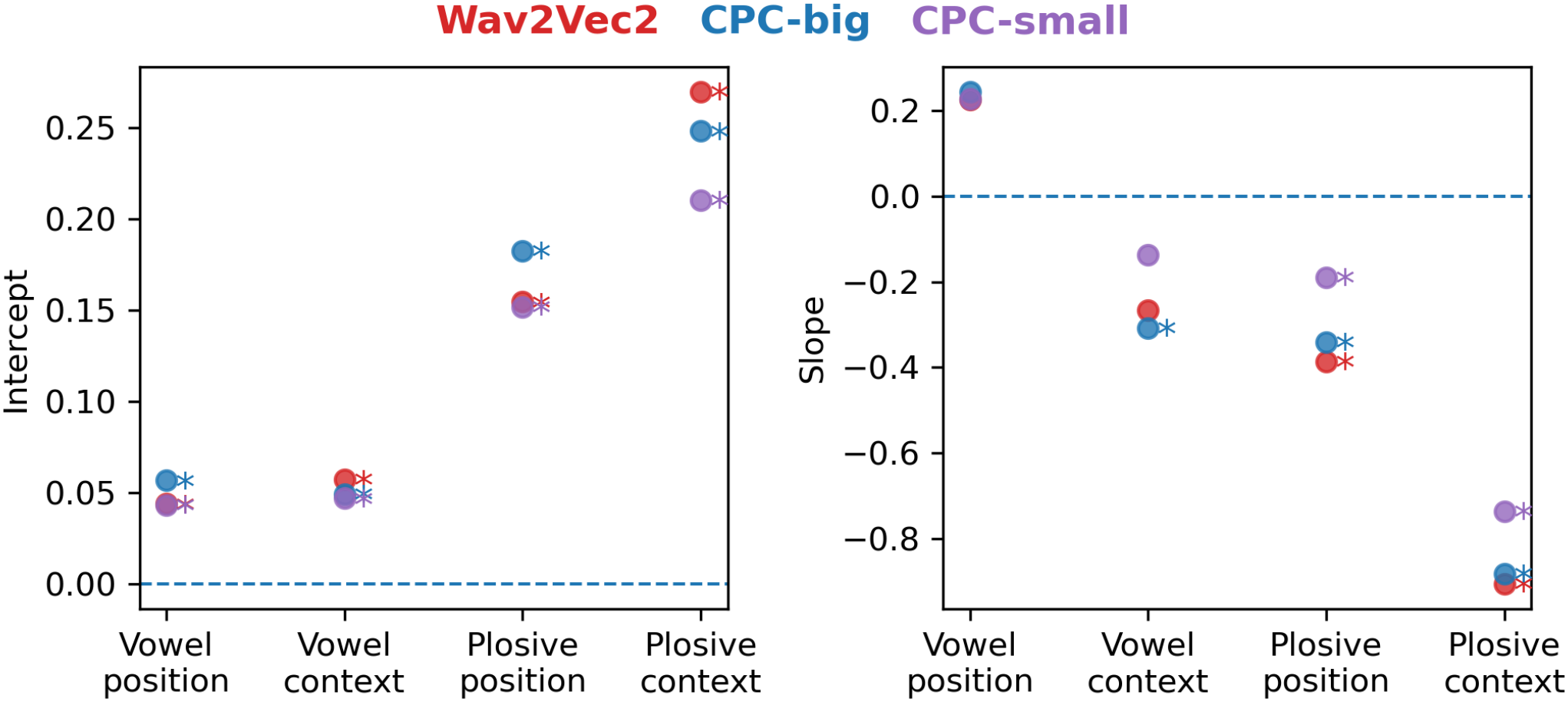
The slope and intercept of ordinary least squares regressor fitted for each speech model (* indicates *p <* 0.05). The predictors in the regression are the mean-centered generalization strengths of acoustic features. The prediction targets are the differences between the generalization strengths of one speech model and that of CPC-audio. All intercepts were estimated to be positive, suggesting all three speech models generalize better than CPC-audio on average in the four cases we considered. The advantage is more pronounced for plosives than for vowels. Slopes were mostly negative except for cross-position generalization of vowel decoders. A negative slope suggests the advantage of the speech model increases as the acoustic generalization decreases, which means the advantage comes from abstracting away from the acoustic variability. This is again most prominent for cross-context generalization of plosives.

## Footnotes

* We say a phone is “encoded” if its identity can be decoded from brain signals. Decodability does not necessarily imply that the information is indeed used in processing.

† Technically, both prediction for CPC and reconstruction for Wav2Vec2 are operationalized as a contrastive training objective computed in the latent space.

‡ Note that the generalization strength can reach over 100%, for example if samples from condition B are similar to, but less noisy than, those from condition A)

